# Somatic disinhibition of granule cells improves information transmission and pattern separation in the dentate gyrus

**DOI:** 10.1101/2023.02.16.528800

**Authors:** Cristian Estarellas, Efrén Álvarez-Salvado, Laura Pérez-Cervera, Claudio R. Mirasso, Santiago Canals

**Affiliations:** Instituto de Física Interdisciplinar y Sistemas Complejos, IFISC (UIB-CSIC), Campus Universitat de les Illes Balears, 07122 Palma de Mallorca, Spain; Instituto de Neurociencias, Consejo Superior de Investigaciones Científicas, Universidad Miguel Hernández, San Juan de Alicante, Spain

## Abstract

Cortical circuits operate in a tight excitation/inhibition balance. This balance is relaxed during learning processes, but neither the mechanism nor its impact on network operations are well understood. In the present study, we combined *in-vivo* and *in-vitro* neuronal recordings with computational modelling and demonstrated that synaptic plasticity in the afferents from the entorhinal cortex (EC) to the dentate gyrus (DG), in addition to strengthening the glutamatergic inputs into granule cells (GCs), depressed perisomatic inhibition. Computational modelling revealed a functional reorganization in the inhibitory network that explained several experimental findings, including depression of the feed-forward inhibition. *In vitro* results confirmed a perisomatic dominance of the inhibitory regulation with important functional consequences. It favoured GCs burst firing, improved reliability of input/output transformations and enhanced separation and transmission of temporal and spatial patterns in the EC-DG-CA3 network.

## Introduction

Excitatory activity recorded in cortical neurons is consistently matched by inhibitory activity of proportional magnitude (Okun & Lampl, 2008). However, inhibition can be modulated independently of excitation, and sustained excitation/inhibition (E/I) imbalance has been reported in periods of sensorimotor integration, increased attention, and the formation of associative memories (Froemke, 2015; Letzkus et al., 2015). Modelling work on the regulation of the E/I balance in neuronal networks has shown the importance of disinhibition in the selective gating of specific signals processing (Kremkow et al., 2010; Vogels & Abbott, 2009). However, the rules and mechanisms that relax the otherwise tight balance between excitation and inhibition are not well understood.

The dentate gyrus (DG) is a structure of the hippocampal formation that is attributed with a function in pattern separation, important for memory formation (Treves et al., 2008). The DG transforms the dense multisensory information that receives from the entorhinal cortex (EC) into a sparse code that minimizes the overlap between the activated neuronal populations, facilitating the discrimination of similar input patterns. The low responsiveness of granule cells (GCs) in the DG circuit is central to the mechanism for pattern separation. Although GCs receive large numbers of simultaneous inputs from the EC (Pernía-Andrade & Jonas, 2014), their low excitability (Schmidt-Hieber et al., 2007) and tight inhibitory control (Ewell & Jones, 2010) result in a low firing activity that filters the EC input (Alme et al., 2010; Pernía-Andrade & Jonas, 2014). Regulation of inhibitory activity in this circuit may therefore play a critical role, as a high inhibitory tone is required for sparse GC coding (Leutgeb et al., 2007), but excessive inhibition would prevent the effective activation of the downstream CA3 network.

The E/I balance in the DG has been shown to depend on different short- and long-term synaptic plasticity mechanisms (Evstratova & Tóth, 2014; Galván et al., 2011; Pelletier & Lacaille, 2008). Already in the pioneering work of Bliss and Lomo (Bliss & Lømo, 1973), in which long-term potentiation (LTP) of synaptic strength was first described, it was documented a shift in the input-output transformation in GCs, so that the same excitatory synaptic input in response to perforant path (PP) stimulation produced a higher firing output after LTP. In experiments combining electrophysiological recordings in the DG with fMRI, the induction of LTP further unveiled a functional reorganization that extended beyond the local DG circuit (Canals et al., 2009). It was shown that LTP in the DG increases the functional coupling within the hippocampus and between the hippocampus and other extrahippocampal structures such as the prefrontal cortex, the perirhinal cortex and the nucleus accumbens (Álvarez-Salvado et al., 2014; Canals et al., 2009; Del Ferraro et al., 2018). Interestingly, this functional reorganization was later replicated by just increasing the E/I balance in the DG by means of inhibiting parvalbumin positive interneurons with pharmacogenetic tools (Caramés et al., 2020). These results suggest that the regulation of GCs firing and information transmission in the EC-DG-CA3 network may be controlled by the inhibitory circuit in the DG and operated by synaptic plasticity.

In this work we combined *in vivo* and *in vitro* neuronal recordings together with computational models (Fig. S1) to investigate the role of synaptic plasticity in the regulation of the E/I balance and communication in the hippocampal network. We found that synaptic plasticity in the DG decouples excitation from inhibition facilitating information encoding and transmission. The key finding was an unexpected reduction of feed-forward inhibition onto GCs driven by synaptic potentiation in the PP, likely produced by the reorganization of the local inhibitory network. LTP-driven disinhibition was found to be predominantly confined to the perisomatic compartment in GCs. This imbalanced disinhibition in the somato-dendritic axis had important functional consequences that we explored in a compartmental neuronal model of the DG (Chavlis et al., 2017). Overall, it favoured GCs burst firing, improved reliability of input/output transformations and enhanced separation and transmission of temporal and spatial patterns in the EC-DG-CA3 network.

## Results

### LTP in the perforant pathway depresses feedforward inhibition in GCs

We performed electrophysiological experiments in anesthetized rats employing linear array electrodes across the dorsal hippocampus and stimulation electrodes in the PP (Fig. 1A, see Supplementary Methods, Fig. S2A (Álvarez-Salvado et al., 2014; Canals et al., 2009)). By PP stimulation (Fig. S2B) and employing Independent Component Analysis (Fig. S2C (Benito et al., 2014; Fernández-Ruiz & Herreras, 2013; O. Herreras et al., 2015; Oscar Herreras, 2016; Łęski et al., 2010; López-Madrona et al., 2020; Makarov et al., 2010; Makarova et al., 2011; Ortuño Silva et al., 2019; Schomburg et al., 2014)) we were able to separate the DG evoked LFP (e-LFP) in two components called e-PP and e-Hilar, according to their topographic localizations (Fig. 1B and Fig. S2B,C). Accordingly, these components present dipoles in the Current Source Density (CSD) depth profile (Fig. 1B) in the mid-molecular layer and the hilus, respectively. More specifically, the e-PP matched the terminal field of the medial entorhinal cortex layer II (MEC), with clear CSD sinks (inward currents) and excitatory postsynaptic potentials (EPSPs) in response to the PP stimulation (López-Madrona et al., 2020). The e-Hilus, centred in the hilus, presented active current sources (outward currents) in response to PP stimulation, located in the granule cell layer (putative perisomatic inhibition). The latency of the evoked potentials was 5.1±0.03 and 7.4±0.05 ms (mean ± sem, n=8 animals) for the e-PP and e-Hilar, respectively, consistent with monosynaptic and disynaptically (feedforward) evoked potentials (Fig. S2C). Administration of GABAergic antagonists, such as bicuculine, gabacine or CGP, by microinjection in the DG, selectively blocked the e-Hilar component without altering the e-PP (Fig. S2D, E, F). Overall, these results demonstrate that a readout of the E/I balance in the DG can be extracted from multichannel LFP recordings and the use of IC analysis.

**Figure 1.**
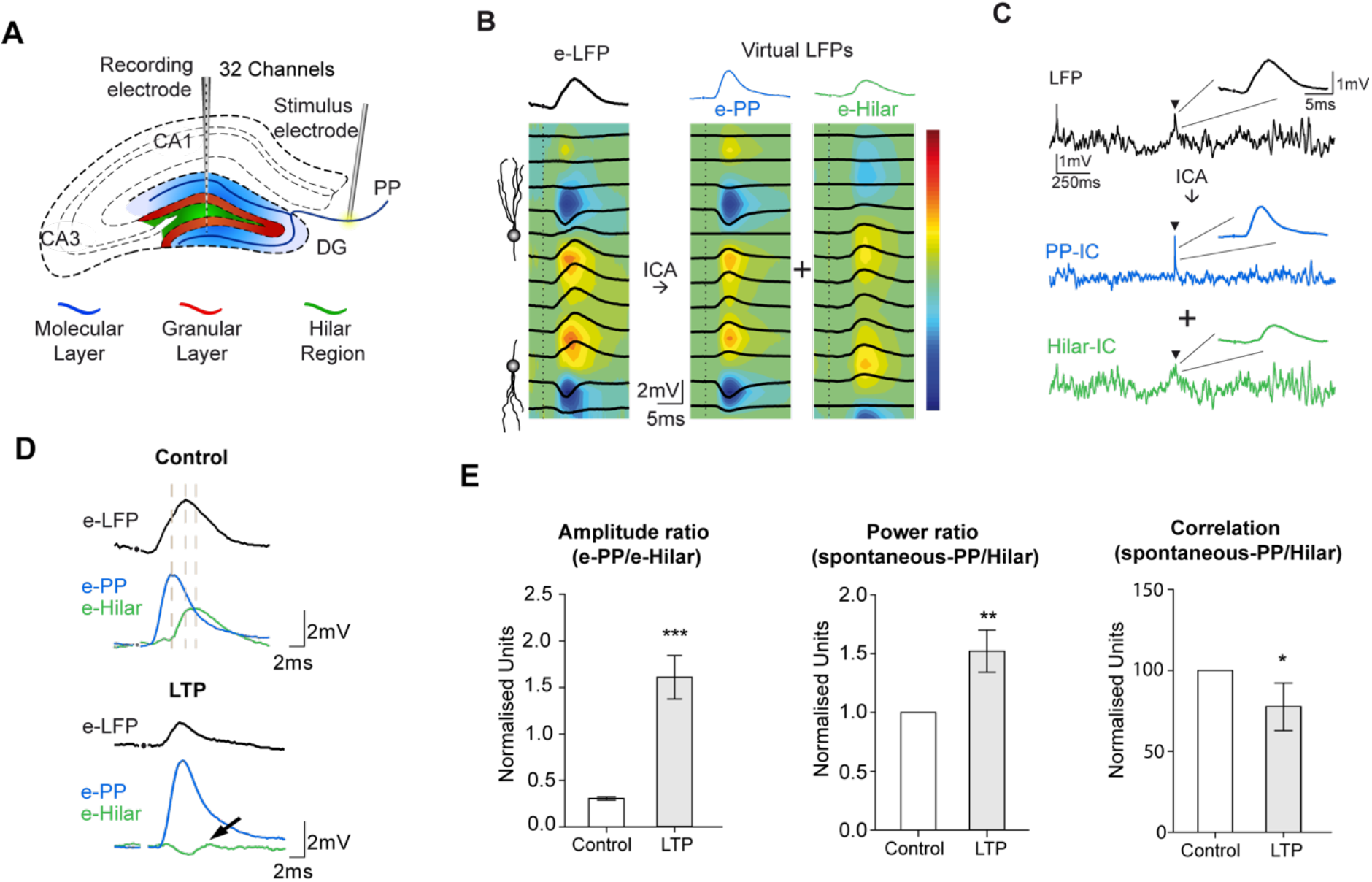
Recording of excitatory and inhibitory components of the DG LFP. **A)** Scheme of the hippocampus highlighting the DG in color (blue: molecular, red: granular, green: hilar layers) and illustrating the location of recording and stimulating electrodes. **B)** Left: Evoked LFP recordings (e-LFP) across the DG in response to a PP stimulation overlaid on the corresponding CSD map (color coded). The upper trace (black) highlights the evoked potential in the center of the hilar region. Right: Decomposed ICs (virtual LFPs) overlaid on the corresponding CSDs are shown for the PP-IC (e-PP, middle panel) and Hilus-IC (e-Hilus, right panel). Upper traces highlight the corresponding e-PP (blue) and e-Hilar (green). **C)** Spontaneous activity in the LFP, PP-IC and Hilus-IC corresponding to the experiment in B. The black arrow indicates the evoked response shown in B. **D)** Comparison of the evoked potentials in the raw LFP and ICs in the control condition and after LTP induction. The arrow points to the evoked potential in the Hilus-IC that is strongly reduced after LTP. **E)** LTP effects on the evoked IC’s amplitude ratio (left, t(23)=6.6, p<0.0001), on the power of the spontaneous IC’s (middle, t(11)=2.9, p<0.01), and their correlation (right, t(11)=2.1, p<0.05), normalized to the average value in each experiment. Bars represent mean ± SEM.

We then used layer-specific LFPs to investigate the effect of synaptic plasticity on the E/I balance. We induced LTP in the PP (see methods) and recorded both PP-evoked potentials and spontaneous activity. The comparison of the evoked subthreshold potentials in each generator before and after LTP induction revealed two distinct effects (Fig. 1D). First, LTP produced the expected potentiation of the glutamatergic input on GCs (i.e. e-PP; fig. 1D, upper traces; see also Fig. S3 for suprathreshold stimulus-response curves) and, second, a strong depression of the feed-forward inhibition recorded in the e-Hilar (Fig. 1D, lower traces). As a consequence, the E/I ratio extracted from evoked components largely increased (Figure 1E, left panel). In the spontaneous activity, we measured the power of each component and the correlation between them (Fig. 1E, middle and right panels respectively), and found that LTP increased the PP/Hilar ratio, and decreased their correlation, indicating that LTP induction decoupled excitatory and inhibitory activities.

### LTP preferentially depressed perisomatic inhibition

The inhibitory circuit of the DG is composed by a wide range of inhibitory interneurons (Houser, 2007). As shown in Fig. 1C, the electric currents in the Hilus-IC depressed by LTP are predominantly located in the perisomatic region in the granular cell layer. To corroborate this finding and discriminate perisomatic from dendritic inhibition, we performed intracellular recordings in an *in vitro* preparation. However, due to the high inhibitory tone in the DG slice preparation, LTP induction requires the use of pharmacological antagonists of GABA receptors (Arima-Yoshida et al., 2011; Chen et al., 2001; Vyleta & Snyder, 2021). This prevents the study of inhibition in such preparation. To overcome this limitation, we chose a combined protocol of *in vivo* LTP induction (or control “sham” stimulation) and *in vitro* recordings (Fig. 2A, see methods).

**Figure 2.**
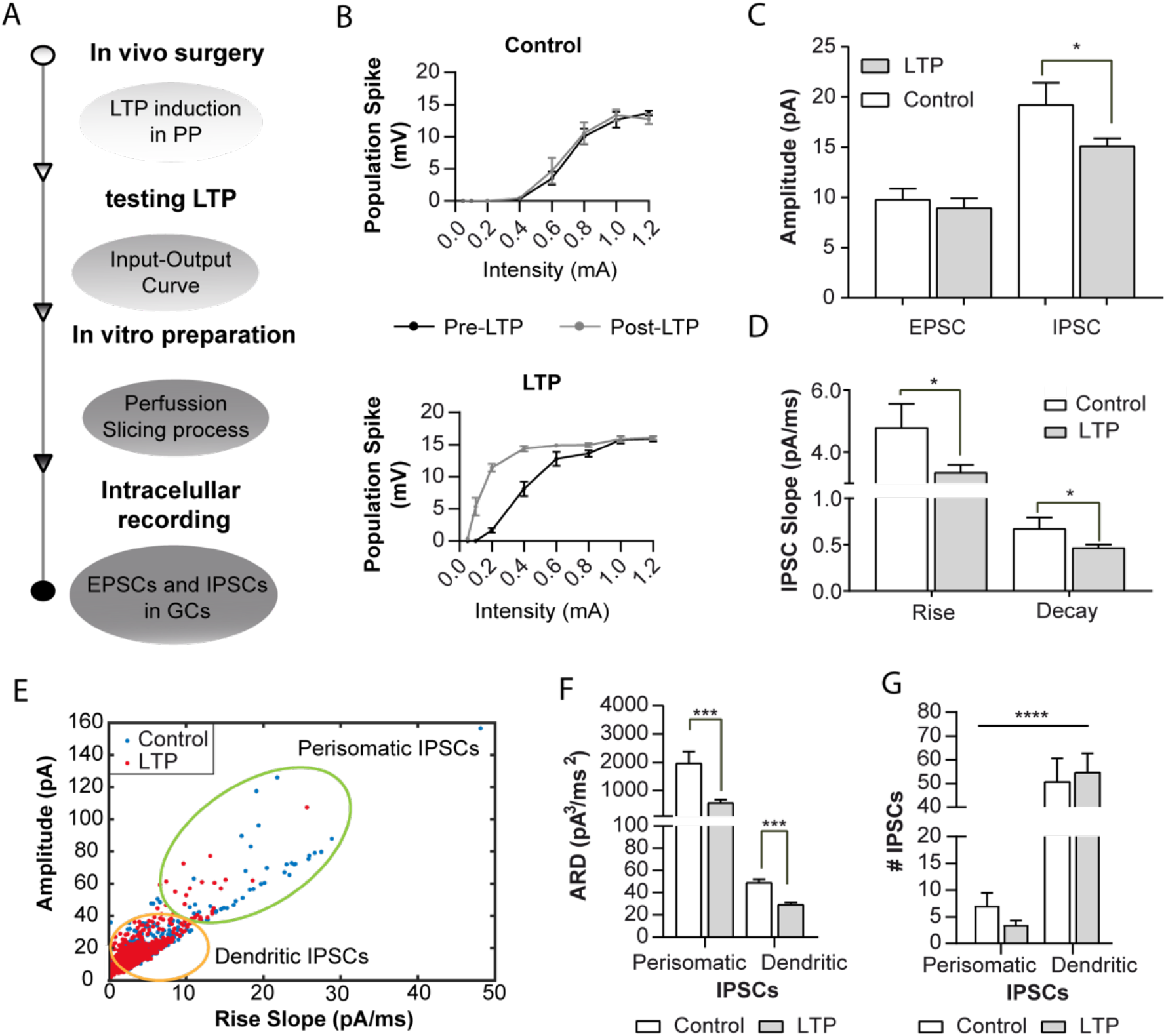
Dendritic and perisomatic inhibitory inputs respond differently to LTP. **A)** Scheme of the combined *in vivo* and *in vitro* experiments. **B)** Representative stimulus-response curves obtained from control and LTP experiments. **C)** Amplitude of the spontaneous EPSCs and IPSCs recorded in GCs (mean ± SEM) (EPSC: one-tailed t(23)=0.6, p=0.3; IPSC: one-tailed t(12.4)=1.8, p=0.048) **D)** Rise and decay slopes of the IPSCs recorded in GCs (mean ± SEM) (Rise: one-tailed U(11,14)=42, p=0.029; Decay: one-tailed t(23)=1.84, p=0.0392). **E)** K-means classification of the inhibitory postsynaptic currents depending on their amplitude, rise slope and decay slope (only represented rise slope and amplitude for visualization purposes). The two populations are schematically identified by the orange (dendritic) and green (perisomatic) circles. **F)** Product of the amplitude, rise and decay slopes (ARD) from perisomatic and dendritic IPSCs recorded in control and LTP groups (Perisomatic: U(93,55)=1684, p=0.0005; Dendritic: U(664,880)=259692, p=0.0002). **G)** Number of perisomatic and dendritic IPSCs per cell (control n=13, LTP n=16) in 100 s (# IPSCs; mean ± SEM). The statistic represents the difference in the distribution of perisomatic and dendritic IPSCs in GCs between Control and LTP cases (Fisher’s exact test, p<0.0001).

After the *in vivo* preparation in which half of the animals received an LTP induction protocol (Fig. 2B), we prepared slices and recorded spontaneous EPSCs and IPSCs in the GCs *in vitro* to measure the state of the local E/I circuit. As shown in Fig. 2C, in control conditions there is a clear dominance of inhibitory *vs.* excitatory currents in the slice, as expected (Kraushaar & Jonas, 2000; Liu et al., 2014; Nitz & McNaughton, 2004; Stefanelli et al., 2016). However, we found a significant reduction in the amplitude (Fig. 2C), as well as in the rise and decay time constants (Fig. 2D), of the IPSCs in LTP cases as compared to controls. These results corroborated the reduction of the inhibitory tone over GCs after LTP induction in the PP *in vivo.* The failure to find *in vitro* an increase in the amplitude of EPSCs in the LTP group is attributed to the slice preparation in which the potentiated pathway originating in the entorhinal cortex was resected.

Inhibitory inputs recorded from the soma reflect their dendritic or perisomatic origin in the amplitude and kinetics of the generated currents, with perisomatic inputs being larger and faster (Rall et al., 1992). We reasoned that a selective reduction in perisomatic inhibition would reduce the variance of the recorded IPSCs, homogenizing the remaining inputs towards lower amplitude and slower kinetics. Indeed, we found decreased variance in the distribution of IPSCs amplitudes after LTP (F-test, p-value = 0.00026). Then, using the k-mean clustering method (Materials and Methods), we classified the IPSCs in dendritic or perisomatic according to their amplitude, rise and decay slopes (Figure 2E). Both perisomatic and dendritic inhibitory inputs were reduced after LTP induction (Fig. 2F), however, the proportion of perisomatic inputs was smaller than those of dendritic origin (Fig. 2G), causing a somatodendritic imbalance over the granule cells.

### LTP activates the hilar circuit responsible for the excitation/inhibition imbalance

To elucidate the mechanism responsible for the E/I imbalance we built a neural circuit of the DG (see Materials & Methods). Five neuronal populations were considered as the representative network of the DG (Amaral et al., 2007; Pelkey et al., 2017): Medial Entorhinal Cortex LII (MEC), Granular cells (GCs), basket cell interneurons (BC), hilar inhibitory interneurons (Hil) and Mossy Cells (MC) (Fig. 3A). Neurons were modelled using the Izhikevich equations (Izhikevich, 2003) and assumed to be connected by chemical synapses simulating the kinetics of the most common receptors participating in the circuit (Destexhe et al., 1995): NMDA, AMPA and GABA_A_. The circuit parameters were tuned to reproduce the observed dynamics of the DG, in particular, the characteristic theta-gamma frequency component recorded *in-vivo* in rats (Pernía-Andrade & Jonas, 2014). The MEC input mainly oscillates in the theta frequency range (Buzsáki, 2002; Mitchell & Ranck, 1980; Stepan et al., 2012) while the Hilus region does it in the gamma band (Bragin et al., 1995; Trimper et al., 2017), as shown in Figure 3B. With the chosen parameter values the model reproduced the coherence between the EPSCs and IPSCs in GCs and the LFP simultaneously recorded in the DG (Pernía-Andrade & Jonas, 2014), as shown in Figure 3C. For simplicity and without losing generality, we modelled LTP as an instantaneous change in the synaptic weights in the PP inputs.

**Figure 3.**
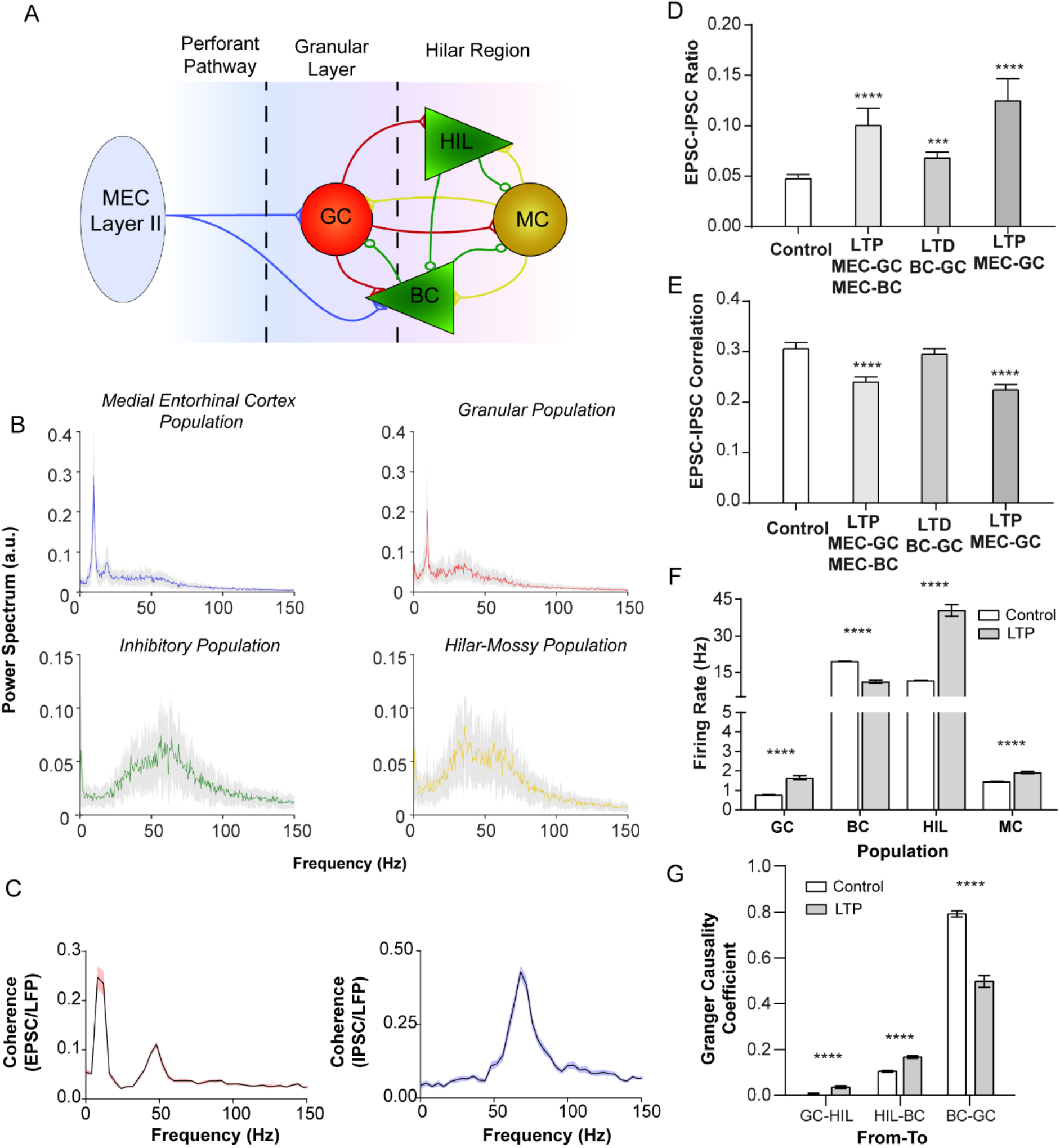
Modelling results of the LTP-induced reorganization of the DG circuit. **A)** Scheme of the DG circuit’s model. Five populations were considered: Medial Entorhinal Cortex (MEC), Granular Cells (GC), Mossy Cells (MC), Basket Cell (BC), and Hilar (HIL) inhibitory interneurons. Circles and triangles at the end of the connections indicate excitatory and inhibitory synapses, respectively. **B)** Power spectra of different cell populations in the model. **C)** Coherence between the granular LFP (average population voltage) and the excitatory (left) and inhibitory (right) postsynaptic currents in individual GCs. **D-E)** Ratio amplitude and correlation between the excitatory and inhibitory activity onto GCs. Different potential effects of LTP are tested and compared to the baseline (control) condition: concomitant potentiation of the MEC connection to GCs and BCs, depression of the BCs connection to GCs, and potentiation of the MEC connection to GCs alone. Ratio (D): H(3)=33.24,p<0.0001; Post-hoc Dunn test: Control vs LTP MEC-GC MEC-BC (p<0.0001), Control vs LTD BC-GC (p=0.03), Control vs LTP MEC-GC (p<0.0001). Correlation (E): H(3)=45.03,p<0.0001; Post-hoc Dunn test: Control vs LTP MEC-GC MEC-BC (p<0.0001), Control vs LTD BC-GC (p>0.9), Control vs LTP MEC-GC (p<0.0001). **F)** Firing rate of the different DG populations (mean ± sem). GC: t(38)=8.3, p<0.0001; BC: t(38)=12.6, p<0.0001; HIL: t(38)=12, p<0.0001; MC: t(38)=7.4, p<0.0001. **G)** Granger Causality between the inhibitory populations and GCs. GC-HIL: t(38)=5.6, p<0.0001; HIL-BC: t(38)=8.6, p<0.0001; BC-GC: t(38)=10.3, p<0.0001.

While the effect of LTP on GCs is well stablished, its effect on interneurons receiving PP inputs, as the BCs, is not well known. Consequently, we simulated different scenarios. A depression of the PP-evoked feed-forward inhibition, as observed experimentally, can be obtained by simply assuming a reduction in the BCsàGCs input. Although this effect yielded an increased excitation/inhibition ratio, it did not change the correlation between the excitatory and inhibitory components as found experimentally *in vivo* (see Fig. 1E). The most parsimonious alternative was, however, to consider an equal potentiation to both MEC projections to GCs and BCs. As a result, we found a reduction of the EPSC/IPSC amplitude ratio together with a smaller correlation between these currents (Fig. 3D and E), in agreement with the experimental findings (Fig. 1). Similar results were obtained by the sole potentiation of the MECàGC connection (Fig. 3D and E).

We next analysed numerically the firing rate of each population in the model (Fig. 3F) and computed the Granger Causality that accounts for the directional flow of information in the circuit (Fig. 3G). Our results suggested that the increased firing of GCs as a consequence of MEC potentiation drives Hil interneurons. In turn, hyperactive Hil interneurons projecting to BC interneurons reduced feed-forward inhibition. It is worth noting that Hil interneurons and the MCs were key components in the circuit, as the removal of any of these two populations deleted the LTP effect (Fig. S4).

### Perisomatic disinhibition improves DG encoding and transfer to CA3

The previous model provided a plausible explanation for the mechanism yielding the experimentally observed E/I imbalance in GCs. Furthermore, *in vitro* experimental findings unveiled a segregation of LTP effects between the perisomatic and dendritic inhibitory compartments. However, what are the functional consequences of this segregation was not known. To address this question, we built a multicompartmental GC model, following the work Chavlis et al., 2017, containing three dendritic branches and the soma (a total of 27 compartments). As shown in Figure 4A, we considered GCs excitatory inputs at medial and proximal dendritic compartments from the MEC and MCs, respectively. Inhibitory inputs from hilar PP associated (HIPP) cells imping on distal dendritic segments, and hilar commissural associational path (HICAP) interneurons innervate proximal segments. BCs were modelled as single compartment units that projected to the soma of the GC and received excitatory inputs from MEC, MCs and GCs and inhibitory inputs from the Hil population (Fig. 4A). Our GC model could generate both regular spiking and bursting regimes (Fig. S5), according to the experimental literature (Mistry et al., 2011; Pan & Stringer, 1996). Through the MEC input we introduced different temporal and spatial patterns and computed the GC output.

**Figure 4.**
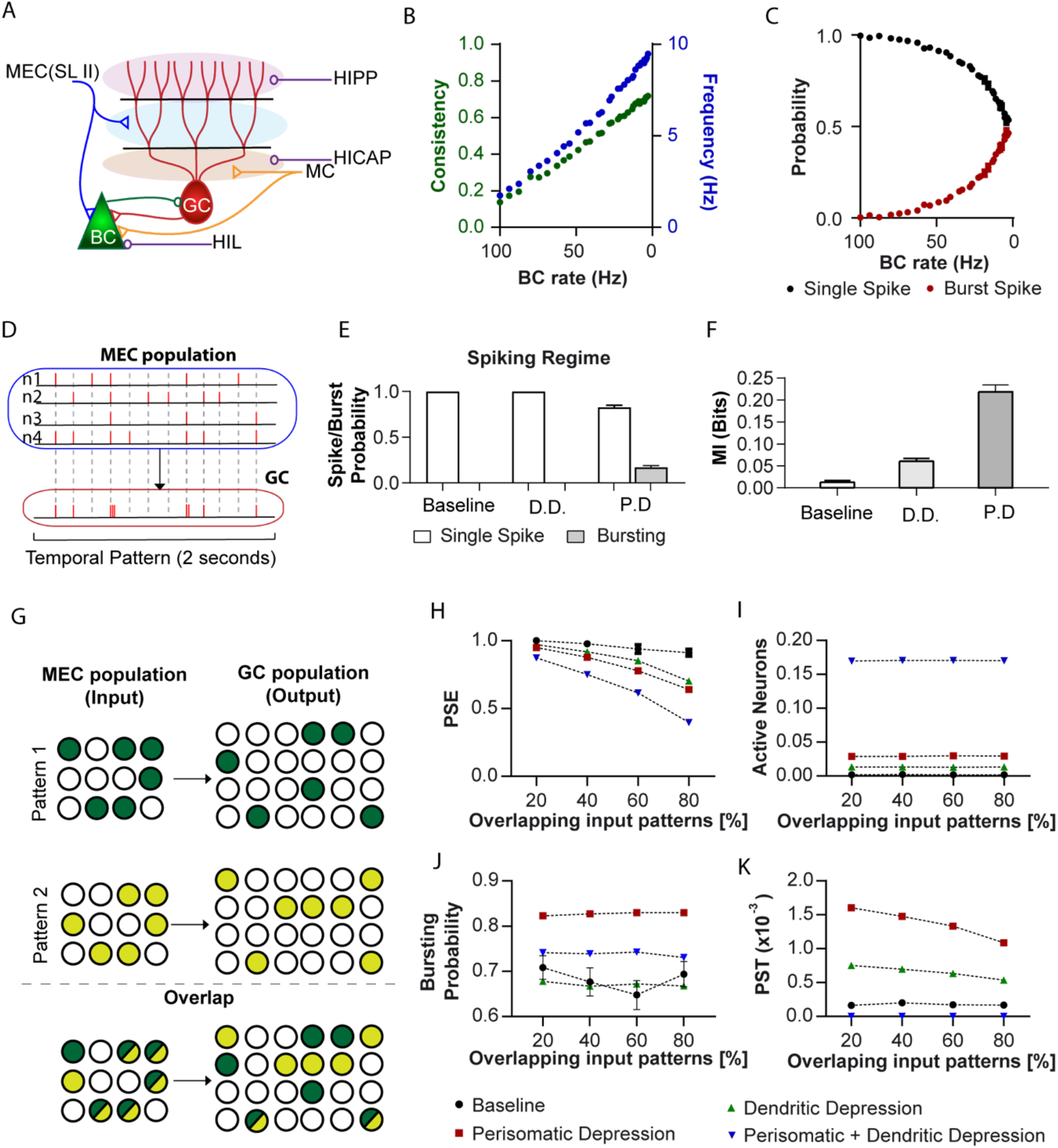
Functional role of inhibitory perisomatic vs. dendritic inhibition over the GCs. **A)** Scheme of the detailed multicompartmental model of an individual GC coupled to different inhibitory and excitatory sources. Circles/triangles indicate inhibitory/excitatory synapses, respectively. MEC, MC and HIL sources represent inputs from the Medial Entorhinal Cortex, Mossy Cell and Hil populations (see Figure 3A). HICAP and HIPP sources represent inhibitory inputs into distal and proximal dendrites. **B)** Consistency and firing frequency of the GC output and **C)** Probability that the GC fires in spikes or bursts, as a function of the BC firing rate. **D)** Scheme of the temporal pattern encoded into the GC. Four patterns of 2 seconds duration each were generated and injected into the GC via the MEC input. The process was repeated for 30 seconds. **E)** The same as **C)** but for the temporal pattern input; Baseline, corresponds to the original configuration of the inhibitory inputs into the GCs; D.D. (P.D.) corresponds to dendritic (perisomatic) depression, i.e., when dendritic (perisomatic) inhibitory inputs over GC were reduced. **F)** Mutual Information between input from MEC and output of the GCs. **G)** Scheme of the spatial Pattern Separation Task (PST). 1000 independent GCs were considered. **H)-K)** Pattern Separation Efficiency (PSE) (**H**), number of activated neurons by the pattern (**J**), probability of generating bursts (**K**), and Pattern Separation Transmission (PST) (**K**) for a pair of input patterns as a function of the percentage of overlap between patterns. All measurements are the mean ± sem over fifty random pairs of input patterns.

We first investigated the contribution of perisomatic inhibition to the GCs input/output transformation. We found that the consistency, measured as the correlation between two GC outputs in response to the same MEC input, increased when decreasing BC activity. In these conditions, GC firing rate and the probability of burst-firing also increased (Fig. 4B, C). Next, we studied the temporal coding capacity of GCs. We generated 4 temporal patterns of MEC activity that impinged simultaneously on GCs (see methods; Fig. 4D), and quantified the Mutual Information (MI) between the MEC input and GC output. We found that disinhibition increases MI, with disinhibition of the perisomatic compartment being more efficient than disinhibition of the dendritic compartment (almost tripling MI, Fig. 4F).

An increase in activity, consistency, and input-output MI in GCs might indicate enhanced temporal encoding. However, a function like pattern separation might suffer from the increased excitation of GCs and the possible reduction in sparseness (Madar et al., 2019; Petrantonakis & Poirazi, 2014, 2015; Treves et al., 2008). To investigate the impact of compartment-specific disinhibition on pattern separation we extended the model by considering 1000 independent GCs and used two spatial MEC input configurations that generated GC activity patterns with different degrees of overlap (Fig. 4G). We started by evaluating the Pattern Separation Efficiency (PSE), defined as the inverse of the normalized overlap between two patterns (see Material & Methods). In Figure 4H, it can be seen that reducing either perisomatic or dendritic inhibition or both of them worsen PSE, as expected from an increase in GC firing rate and the loss of sparseness. However, PSE performance does not necessarily imply a robust transmission of the separated patterns. In fact, the best PSE performance obtained under baseline inhibitory conditions is achieved at the cost of activating a very low number of GCs (Fig. 4I). This occurred due to the tight inhibitory control, which resulted in a drastic loss of MI between the MEC input and GC output (Fig. 4F). A balance is then required between sparse firing (to filter the dense EC input), and sufficient mass action to impact the dynamics of the downstream CA3 network. Therefore, a better indicator should consider not only the PSE but also the number of active neurons (AN) (Fig. 4I) and the probability of bursts (PB) firing (Fig. 4J). The latter is known to act as a detonator with a strong impact on CA3 pyramidal cell firing and synaptic plasticity (Vandael et al., 2021). To quantify the balance of population activity on information transmission and pattern separation, we defined the Pattern Separation Transmission (PST) index as a function of the three previous variables (see Methods).

In Figure 4K it can be seen that the PST is largely enhanced when reducing perisomatic inhibition. Dendritic disinhibition also enhanced PST, but to a smaller degree, as it produced a smaller increase in spiking activity and, most importantly, did not contribute to burst firing. When complete depression of inhibition in the perisomatic-dendritic axis is considered, PST is reduced to 0. Therefore, the dominant reduction in perisomatic inhibition induced by synaptic plasticity in the PP might balance firing sparseness and bursting to support pattern separation while optimizing information transmission between the MEC and CA3.

## Discussion

In this work we found, combining *in vivo* and *in vitro* experimental data as well as modelling results, that synaptic plasticity in the perforant pathway, the main input from the entorhinal cortex to the hippocampus, decouples excitation an inhibition facilitating information encoding and transmission. The key finding was an unexpected reduction of feed-forward inhibition onto GCs driven by synaptic potentiation in the PP. Our experiments revealed that this effect predominantly occurs in the GC perisomatic inhibitory compartment, pointing to a main role of BCs. A first computational model of the DG, fitted to experimental data, showed that this disinhibition is likely the consequence of a functional reorganization of the local hilar network. A second model comparing perisomatic and dendritic inhibition highlighted the functional importance of this inhibitory balance. Perisomatic disinhibition improved transmission from the MEC to CA3 (increased MI), input/output transformations became more reliable (consistency), and GCs better transmitted temporal and spatial patterns to CA3 while preserving their pattern separation ability.

Our experimental results showed that LTP regulates both excitatory and inhibitory inputs into GCs. In addition to the well-known potentiation of the glutamatergic input from the EC, the disynaptic feedforward inhibition was depressed, a change that alters the E/I balance and thus the input/output transformations in CGs, in favour of greater responsiveness. The classic work by Bliss and Lomo (Bliss & Lømo, 1973) on the LTP phenomenon showed that firing activity in the population was larger than would correspond to the sole increase in synaptic activity after LTP induction. The so called EPSP-to-spike potentiation could be in part the result of a reduced feed-forward inhibition as the one shown here. This mechanism may also explain the results obtained in brain imaging experiments (fMRI) of LTP showing increased activity propagations within the hippocampus and from the hippocampus to cortical and subcortical mesolimbic structures (Álvarez-Salvado et al., 2014; Canals et al., 2009; Del Ferraro et al., 2018). Our results point to a predominant perisomatic disinhibition in response to LTP, likely involving reduced activity of BCs. Importantly, in a recent work using parvalbumin (PV)-cre mice (Caramés et al., 2020) it is shown that a reduction in the activity of PV+ cells (putative BCs) by means of pharmacogenetic inhibition, mimics the functional reorganization in fMRI networks observed after LTP in the DG in rats, and enhances memory formation in a novel object location task. Overall, our results point to a DG mechanism operated by synaptic plasticity and controlled by BCs that regulates communication in memory networks.

Granule cells of the DG have very low firing rates (Alme et al., 2010; Jung & McNaughton, 1993; Pernía-Andrade & Jonas, 2014) likely due to their low excitability (Schmidt-Hieber et al., 2007) and tight inhibitory control (Ewell & Jones, 2010). Nevertheless, when GCs fire, they do it in bursts with a higher-than-expected probability compared to other principal cells (Pernía-Andrade & Jonas, 2014). Although the mechanism that switches from regular to burst firing in GCs has not been elucidated experimentally, our computational results suggest that perisomatic disinhibition could be a determinant factor. The possibility to switch between regular and burst firing is functionally relevant since the latter acts as detonators enhancing communication between DG and CA3 (Chamberland et al., 2018; Henze et al., 2002; Salin et al., 1996; Vyleta et al., 2016; Zucca et al., 2017). The CA3 response to GC bursts is different in pyramidal neurons than in interneurons (Lituma et al., 2021; Toth et al., 2000). While the response of the former is facilitated, the latter are unchanged or depressed, resulting in an increased excitation/inhibition ratio in CA3. Regulation of burst firing in GCs may represent a mechanism to boost information transmission.

The cognitive consequences of the above mechanism might be reflected in the ability of individuals to differentiate between similar input patterns. DG has been repeatedly linked to pattern separation, for which sparse firing of GCs is considered key (Leutgeb et al., 2007; Marr et al., 1991; Treves & Rolls, 1992). The proposed role of pattern separation in the DG is to prevent that the attractor dynamics of the CA3 network, so appropriate for pattern completion, fuses similar but distinct patterns. It has been shown that when rodents are exposed to very similar environments, their ability to discriminate between them depends on the integrity of the DG (Goodrich-Hunsaker et al., 2008; Hunsaker & Kesner, 2013; Morris et al., 2012). Our results show that decreased inhibition in the DG may increase the overlap between the patterns represented by GCs and compromise pattern separation locally. However, we argue that to be an efficient pattern separator, the orthogonalized output of the DG needs to reach the CA3 region with sufficient strength to push local dynamics away from stablished attractors. Supporting this view, our results show that if the segregation of patterns is solely based on the sparse firing of a low number of GCs, the ability to transmit them to CA3 is compromised. Conversely, a certain level of overlap in the DG may be acceptable if the higher proportion of GCs used to represent each pattern ensures transmission to CA3, for which bursting GCs are particularly suitable. Therefore, a balance between pattern separation and transmission is necessary. Our results indicate that perisomatic disinhibition optimally balances both properties, retaining the ability to separate patterns and markedly increasing the ability to efficiently transmit them to CA3. Indeed, we have recently shown that a selective pharmacogenetic inhibition of BCs in the DG improved pattern separation (Caramés et al., 2020).

In conclusion, we proved by experimental and computational means that synaptic potentiation in the perforant pathway connecting the entorhinal cortex with the DG not only facilitates excitatory glutamatergic activity in GCs, but specifically decreases perisomatic feedforward inhibition involving BCs. This has important functional consequences. The fine tuning between perisomatic and dendritic inhibition controlled by synaptic potentiation balances firing sparseness and bursting, enhancing consistency of GC outputs, and, as a consequence, supporting pattern separation by improving information transmission between the MEC and CA3.

## Acknowledgements

The authors acknowledge funding from the Spanish Ministerio de Ciencia e Innovación, Agencia Estatal de Investigación (PID2021-128158NB-C21, PID2021-128158NB-C22 /10.13039/ 501100011033) and Programs for Centres of Excellence in R&D Severo Ochoa (CEX2021-001165-S /10.13039/ 501100011033) and María de Maeztu (CEX2021-001164-M /10.13039/501100011033). S.C. was further funded by the Generalitat Valenciana Government through the Prometeo Program (PROMETEO/2019/015).

## Methods (Online)

### Animals

Male Sprague-Dawley rats were used for *in vivo* LTP experiments, with a weigh of 250-300 g. Adult C57BL/6J male mice (28-30 g) were used for the combined *in vivo* + *in vitro* experiments. All animals’ procedures were approved by the Animal Care and Use Committee of the Instituto de Neurociencias de Alicante (Alicante, Spain) and comply with the Spanish law (53/2013) and European regulations (EU directive 2010/63/EU).

### Drugs

To inhibit GABA_A_ receptors we used Gabacine 1 mM (SR95531 hydrobromide, GABA_A_ type receptor-specific antagonist; Tocris Bioscience, Bristol, UK), and Bicuculine 100 μM (Bicuculine methiodide, GABA_A_ type receptor-specific antagonist; Sigma-Aldrich, Missouri, USA). To inhibit GABA_B_ receptors, we used CGP 1mM (CGP 52432, GABA_B_ type receptor-specific antagonist; Tocris Bioscience, Bristol, UK).

### *In vivo* electrophysiology surgery

Rats were anaesthetized with 1.2–1.5 g/kg of urethane (Sigma-Aldrich, Missouri, USA) injected intraperitoneally. Supplemental doses (10% of the initial dose) were applied when required. After confirming the absence of reflexes, animals were placed in a stereotaxic frame (Narishige, Tokyo, Japan). During the experiment, the temperature was kept at 37°C, and blood oxygen saturation, heart and breathing rate monitored. After a subcutaneous dose injection of 8 mg/kg of local anaesthetic (Bupivacaine, Braun Medical SA, Barcelona, Spain), the scalp and periosteum were separated. The skull was opened with a manual drill (2 mm diameter, Fine Surgery Tools, USA) over the dorsal hippocampus (coordinates with respect to Bregma: A-P −3.5 mm, M-L 2.6 mm, 3.2–3.5 mm ventral to the dural surface) and the medial Perforant Pathway (PP; coordinates with respect to lambda: A-P 0 mm, M-L 4.1 mm, 2.3–2.7 mm ventral to the dural surface, with an angle of 15° at the sagittal plane directing the tip to rostral) (Paxinos & Watson, 2007). The dura was carefully punctured at the craniotomies with a needle, making the smallest hole possible to facilitate the penetration of both electrodes.

Mice were anaesthetized with isofluorane (Laboratioris Esteve, Murcia, Spain) 4%, 0.8 L/min oxygen for induction, and 1-2%, 0.8 L/min oxygen for maintenance. After confirming the absence of reflexes, animals were placed in a stereotaxic frame (Narishige, Tokyo, Japan) and the scalp and periosteum were separated. During the experiment, the temperature was kept at 37°C and blood oxygen saturation, heart and breathing rate monitored. The recording and stimulating electrodes were implanted following the stereotaxic standard procedures in the dorsal hippocampus (coordinates with respect to Bregma: −2 AP, +1.5 ML, −2 DV) and in the MPP (coordinates with respect to Bregma: −4.3 AP, +2.5 ML, +1.4 DV, - 12° at the sagittal plane).

For orthodromic stimulation of the dentate gyrus, a tungsten bipolar electrode (10-15 kΩ, 325 μm diameter, World Precision Instruments, Florida, USA) was positioned in the medial PP. A multielectrode silicon probe (32 recording sites, 100 and 50 μm inter-site distance, for rats and mice respectively, 413 μm^2^ electrode area, Neuronexus Technologies, Michigan, USA) was placed at the dorsal hippocampus to record the LFP. An Ag/AgCl wire (World Precision Instruments, Florida, USA) electrode was placed in contact with the skin bounded surgery area and used as ground. We found the accurate position of both electrodes using as a reference the control evoked potentials at the dentate gyrus (Andersen et al., 1966) obtaining maximal population spike in the dentate gyrus. The brain electrophysiological signals were filtered (high-pass 0.1 Hz), amplified and digitalized (20 kHz acquisition rate) (Multi Channel Systems, Reutlingen, Germany), and stored for posterior analysis.

### *In vivo* pharmacological experiments

In some experiments, in addition to the stimulating and recording electrodes, a borosilicate glass pipette (World Precision Instruments, Florida, USA) was implanted to deliver different pharmacological agents (dissolved in artificial cerebrospinal fluid). These pipettes were further equipped with Ag/AgCl electrodes to guide their precise implantation in the hilus close to the multielectrode probe. The pipette tip was bent at an angle of approximately 90° to achieve closer proximity to the recording electrode. In all cases, the drug was released through the pipette with an air puff delivered with a picospritzer (custom-built), except for the Gabacine, that was administered by microiontophoresis. Control experiments with artificial cerebrospinal fluid demonstrated the absence of non-desired electrophysiological changes due to volume injection in the above conditions.

The effect of the administered drugs was evaluated measuring the evoked potential in the DG in response to a single pulse stimulation in the PP subthreshold for eliciting population firing. This stimulation facilitates the separation of the pathway-specific evoked potentials (see below: Makarov et al., 2010; Makarova et al., 2011). The only exception was CGP, which effect was quantified in the reversion of the effect of a pair-pulse protocol in which the first pulse is suprathreshold and the second, 150 ms later, is subthreshold. In this protocol, the first pulse conditions the response of the second, producing a GABA_B_-mediated presynaptic inhibition of the GABAergic transmission (Davies et al., 1991; Mott & Lewis, 1991; Vogt & Nicoll, 1999). The protocol was used to demonstrate the GABAergic nature of the evoked hilar potential (e-Hilar).

### *In vivo* LTP protocol

In rats, LTP was regularly induced using a high-frequency stimulation protocol of the PP (Davis et al., 2000). This tetanic stimulation consisted of six trains of pulses (400 Hz, lasting 20 ms), delivered at a 10 s interval, and repeated six times at an interval of 2 minutes. In mice, LTP was induced by a standard theta burst protocol (six 400 Hz-trains of pulses delivered at a 200 ms interval, repeated six times at an interval of 20 s).

To evaluate the synaptic potentiation we measured the population spike, (PS, as the amplitude from the precedent positive crest to the negative peak in the hilar evoked LFP), the excitatory postsynaptic potential (EPSP, as the maximal negative slope of the falling potential in the molecular layer evoked LFP), and the PS latency (as the delay between the stimulation artefact and the PS) at different stimulation intensities (the called input-output curve). We collected Input-Output curves before and after (30-60 minutes) the tetanizing protocol.

### Combined protocol: *in vivo* + *in vitro* electrophysiology

After the *in vivo* induction of LTP (explained above) we perfused the animal with cold NMDG solution followed by rapid decapitation. The brain was dissected and 350 μm transversal hippocampal acute brain slices prepared on the vibratome (VT1200S, Leica Biosystems Nussloch, Germany) with ice-cold NMDG solution. The slices were transferred to a recovery chamber with NMDG solution at 35°C (bubbled with 95% O2 and 5% CO2) for 30 min and then moved to HEPES solution at room temperature for 2 hours.

Slices were kept in ACSF at room temperature for at least 20 min before they were transferred to the recording chamber with continuous ACSF perfusion (bubbled with 95%O2 and 5% CO2). All recordings were done at 32-34 °C controlled by an automatic temperature controller (TC-324C, Warner Instrument, LLC, Holliston, USA). For the Whole-Cell Patch-Clamp recording borosilicate glass pipettes were used (outer diameter, 1.2 mm; inner diameter, 0.86 mm) with 4-6 MW and filled with Cs^-^ intracellular solution.

Cells were visualized under DM6000FS Microscope (Leica) with the ZylasCMOS camera (ANDOR). For voltage-clamps recording, cells were held at 0 mV for GABAergic postsynaptic recordings and −75 mV for glutamatergic postsynaptic recordings. Data were sampled at 10 kHz and filtered at 2.4 kHz using Multiclamp 700B and Digidata 1440 interfaces (Molecular Devices, Sunnyvale, CA, USA).

### *In vitro* Solutions

NMDG-ACSF (in mM): 92 NMDG, 30 NaHCO_3_, 1.25 Na_2_PO_4_, 20 NaHEPES, 2.5 KCl, 25 Glucose, 10 MgCl_2_(6H_2_O), 5 NaAscorbate, 3 NaPyruvate, 2 Thiourea (ph 7.4 300-305 Osm). HEPES (in mM): 92 NaCl, 30 NaHCO_3_, 1.25 Na_2_PO_4_, 20 NaHEPES, 2.5 KCl, 25 Glucose, 2 CaCl_2_, 2 MgCl_2_(6H_2_O), 5 NaAcorbate, 3 NaPyruvate, 2 Thiourea (ph 7.4 300-305 Osm). ACSF (in mM): 119 NaCl, 24 NaHCO_3_, 12.5 Glucose, 2.5 KCl, 1.25 NaH_2_PO_4_, 1 MgCl_2_(6H_2_O), 2 CaCl (ph 7.4 300-305 Osm). Cs-Intracellular Solution (in mM): 130 CsMeSO_3_, 0.5 EGTA, 4 MgCl_2_, 10 CsCl, 10 HEPES, 5 QX-314 Bromide, 2 MgATP, 10 Na_2_-phosphocreatine, 0.3 NaGTP (ph 7.2 288 Osm).

### Computational Dentate Gyrus Circuit Model (*Sup1.-Computational model 1*)

We used individual spiking neurons governed by Izhikevich equations to build the circuit (Izhikevich, 2003). By modifying the Izhikevich parameters (a, b, c, and d) different spiking regimes can be generated (see Table 1). Fast-spiking parameters were used for inhibitory interneurons and excitatory MCs, while tonic spike neurons were used for the excitatory cells of the EC population and 35% of the granular population. However, 65% of GCs were represented by bursting neurons. (Pernía-Andrade & Jonas, 2014).

**Table 1.** Parameters for simulation of neurons in the *computational model 1*. Value of Izhikevich parameters *(a, b, c, d)* obtained from the study Izhikevich, 2003 to generate the spiking dynamics of the main neurons used in the DG circuit model: Fast-spiking, Regular Tonic Spiking and Bursting neurons.

| Izhikevich parameter (row)/<br>Neuron type (column) | a | b | c | d |
| --- | --- | --- | --- | --- |
| Fast-spiking | 0.1 | 0.26 | -65 | 1 |
| Regular tonic Spike | 0.02 | 0.2 | -60 | 8 |
| Burst Spike | 0.02 | 0.2 | -50 | 2 |

We scaled the DG circuit 1:100 to match the anatomical number of GCs in adult rats, and then simulated 100 BCs, 100 Hil interneurons, and 300 MCs. To reduce the computational cost, we only included the active cells in the granular group, which represented 5% of the whole population (Erwin et al., 2020). Additionally, we simulated the EC population, which consisted of 200 neurons, with 20% inhibitory interneurons and 80% excitatory cells, to generate an input theta into the DG circuit. The total number of cells in the simulation was 1200, allowing for a low computational cost necessary for the fitting process method. The synaptic current was modeled as an ohmic conductance multiplied by the ion channel kinetics and the driving force resulting from the difference in voltage between the postsynaptic and reversal potential. We assumed that the opened ionic channel is instantaneous, simulating only the decay exponential of the closing channel, and differentiated the ionic channels kinetics by the exponential time constant characterized by each synaptic type (glutamatergic and GABAergic synapses, including AMPA, NMDA, and GABA_A_.

To fit the model, we relied on three experimental observations: the theta frequency range of the MEC region (Buzsáki, 2002; Mitchell & Ranck, 1980; Stepan et al., 2012), the gamma frequency range of the Hilus region (Bragin et al., 1995; Trimper et al., 2017), and the excitatory and inhibitory inputs characterized by the theta and gamma frequency components, respectively, from the EC and Hilar regions into the granular population, matching the in-vivo intracellular recordings Jonas’ group (Pernía-Andrade & Jonas, 2014). We modified the connectivity and current intensity parameters for each synapse based on the maximum released neurotransmitters (see TABLE 2). The modifications were made manually for each simulation, and we tested the three experimental observations with the power spectrum of the LFP and coherence between EPSC and IPSC of individual GC with the LFP of the granular population. The model also included a stochastic contribution where each neuron received a glutamatergic noise with a Poisson distribution. Figure 2.B and 2.C show the results. In addition, a Poisson distribution was used to simulate glutamatergic noise, which contributed to the model’s stochastic behavior.

**Table 2.** Parameters of connectivity for the *computational model 1*. The connectivity of a circuit indicates the proportion of randomly selected neurons in the target population that receive input from a single presynaptic neuron in the projecting population. The maximum probability released represents the strength of the chemical synapse between the presynaptic neurons (rows) and the postsynaptic neurons (columns).

|  | Connectivity (%) |  |  |  | Maximum Probability Released |  |  |  |
| --- | --- | --- | --- | --- | --- | --- | --- | --- |
| From (row)/to (column) | GC | BC | HIL | MC | GC | BC | HIL | MC |
| EC | 5 | 2 | - | - | 0.05 | 0.01 | - | - |
| GC | - | 2 | 2 | 2 | - | 0.02 | 0.1 | 0.01<sub>(AMPA)</sub><br>0.03<sub>(NMDA)</sub> |
| BC | 12 | 10 | - | 20 | 0.4 | 0.2 | - | 0.3 |
| HIL | - | 60 | 30 | 20 | - | 0.11 | 0.3 | 0.4 |
| MC | 10 | 20 | 20 | 15 | 0.3 | 0.2 | 0.4 | 0.4 |

### Computational Granular Cell Model (*Computational model 2*)

The simulated circuit for the functional model included GCs and BCs, which were adapted from published scripts. (Chavlis et al., 2017). On the other hand, the remaining cells in the DG were incorporated into the model as synaptic inputs onto the simulated neurons. The BC was implemented using a somatic one-compartment adaptive exponential integrate-and-fire (aEIF) model (Badel et al., 2008), which was fitted to produce a fast-spiking regime. The GC in the model was divided into three dendritic branches with a total of 21 passive slots, modeled by the leaky Integrate-and-Fire equation (lI&F) (Burkitt, 2006). Additionally, a variation of the lI&F model (Burkitt, 2006) was used to simulate the granular somatic compartment, including a second differential equation to add an adaptive variable for generating action potentials. Other DG cells were included in the model as synaptic inputs to the simulated neurons.

The model includes both glutamatergic and GABAergic connections, with AMPA and NMDA receptors mediating the former and GABA_A_ receptors mediating the latter. The synaptic conductance was modelled to include both the rise and decay exponential phases that characterize neurotransmitter transmission. In the case of NMDA receptors, their conductance was also modulated by the voltage of the postsynaptic neuron, as these receptors are blocked by a positively charged magnesium ion that is sensitive to voltage changes. This effect was modelled using a sigmoidal function (Jahr & Stevens, 1990) that multiplied the synaptic conductance.

The GC received GABAergic inputs from HIPP and HICAP interneurons onto the distal and proximal dendrites, respectively, and perisomatic innervation from BCs. Glutamatergic inputs were focused on the proximal dendrite due to commissural AMPA synapses and the medial dendrite representing AMPA and NMDA connections from the MEC. These two glutamatergic sources affect the BCs as well, with an inhibitory contribution from Hilar interneurons. The dendritic inputs, except the MEC, were Poisson spike trains with a gamma firing rate. There were one HIPP and HICAP interneuron and five MCs per GC, maintaining the anatomical distribution of the DG. The BCs received the same MC input. The GABAergic input was constituted by several Poisson spike trains with a gamma firing rate that was modified to change the inhibitory input from the Hilar region. The MEC input in both neurons, GC and BC, was a regular input of 4 neurons with a theta frequency of 8 Hz.

### Computational simulations

The DG circuit model was compiled and executed in FORTRAN90, and the functional model was simulated in BRIAN2 (Python 3.1), both computed in the Nuredduna computer system of the Institute of Cross-disciplinary Physics and Complex Sytems (IFISC).

### Data analysis

All analysis done in this work was performed with the software MATLAB (The Mathworks Inc., Massachusetts, USA), Spike2 (Cambridge Electronic Design Ltd., Cambridge, UK), and Clampfit 10.7 (Molecular Devices, LCC, California, USA).

#### Experimental data analysis

The *in vivo* experiments in rats analyzed LFP signals with minimal cortical activity and stable Theta-band oscillation (4-12 Hz) in CA1. The Independent components (IC) were obtained using the *runica* algorithm of the ICA method, which is part of the EEGLAB MATLAB toolbox (Delorme & Makeig, 2004). The method has been developed and tested using brain signals and numerical models (Makarov et al., 2010; Makarova et al., 2011) and has been validated in previous reports (Benito et al., 2014; Korovaichuk et al., 2010).

The cross-correlation between spontaneous signals was calculated using a temporal window of 500 ms, and the power of the signal was computed as the square of the signal. It was calculated for specific frequency bands including Delta (0.5-4 Hz), Theta (4-12 Hz), Beta (12-30 Hz) and Gamma (30-100 Hz), by integrating the area under the periodogram of the signal in that frequency range using the trapezoidal rule. The sub-threshold amplitude is the difference between the maximum potential peak and the baseline voltage in the evoked potential data, and the latency was measured as the time between the stimulation artefact and the peak of the evoked potential.

In the in vitro experiments, recordings of spontaneous activity were filtered with a Gaussian filter cutoff of 100 Hz. The spontaneous postsynaptic currents were detected using templates with amplitude, rise and decay slopes as parameters, automatically obtained. The K-mean clustering method in MATLAB was used to classify dendritic and perisomatic inhibitory inputs by clustering amplitude, rise slope, and decay slope of all inhibitory events, with the optimal number of clusters determined using the silhouette method.

#### Computational data analysis

Granger Causality is a statistical method used to identify the directionality of causation between two time series, by comparing the ability of past values of one series to predict future values of the other series, with and without the inclusion of past values of the second series. In the computational circuit (Sup1. Computational model 1), the Excitatory and Inhibitory currents over the GCs were normalized and analysed as the spontaneous *in vivo* recordings. The total signal was discretized in temporal windows with high overlap between them to improve the analysis accuracy for coherence analysis. The MVGC Multivariate Granger Causality Matlab Toolbox (Barnett & Seth, 2014) is a software package that provides a set of tools for Granger causality analysis of multivariate time series. In the computational circuit described, the Granger causality was used to determine the directionality of causation between the average voltage of each population of cells.

The functional model (Sup1. Computational model 2) focused on studying the information processing of an individual GC model and the pattern separation function in a granular population model. For the individual granule cell, a regular synchronized signal of 4 neurons with a theta frequency of 8 Hz was used as input from MEC in GC and BC. The consistency was evaluated by measuring the spike correlation of the granular output between each non-repetitive combination pair of interactions across 50 iterations. In the spatial codification simulation, each MEC neuron was assigned a firing probability with a theta frequency, generating a repeating pattern of two seconds for 30 seconds. Mutual Information (MI) (Shannon, 1948) was measured for two discrete binary variables - the spike activity of input from MEC and the GC output, represented by 1-spike and 0-no spike.

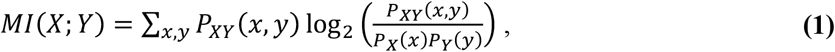

where, X and Y are the discrete variables, *P_XY_* the joint probability distribution *P_X_*(*x*) and *P_Y_*(*y*) are the marginal probabilities.

For the Pattern Separation function, the circuit was scaled up to include 1000 GCs, with the number of neurons in each population determined based on the proportion found in the granular layer: 200 MEC neurons, 50 BCs, 40 MCs, 20 HIPPs, and 20 HICAPs. In this case, the MEC input consisted of a neural pattern of synchronized neurons, constituting 40% of the total MEC population and repeated at a theta frequency for 30 seconds. For further analysis, we selected the neurons that were activated in more than 80% of the input patterns. The Pattern Separation Efficiency (PSE) was calculated by subtracting the ratio of common neurons between a pair of two patterns from the maximum value of 1 (PSE_MAX_).

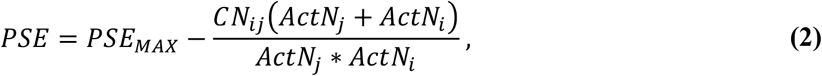

where *CN_ij_* is the number of common neurons in the pair of patterns *i.* and *j,* and *ActN* is the number of active numbers in a given pattern. We also defined a Pattern Separation Transmission (PST) index as:

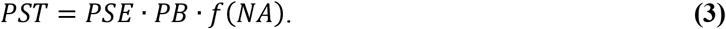

For the function *f*(*NA*) we considered two limits’ cases yielding zero transmission: NA=0 and population overexcitation threshold. A simple equation that allowed us to represent this effect is of the form:

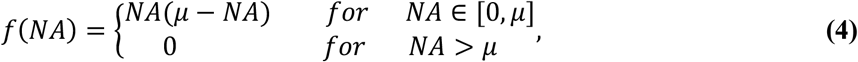

where the parameter *μ* controls the overexcitation of the system. Its value was determined by assuming that the number of active neurons NA that maximizes *f*(*NA*), i.e., that provides maximum information transmission, corresponds to the experimentally estimated number of active granule cells NA=5 % (Danielson et al., 2016; Erwin et al., 2020) i.e., *μ* ≅ 0.1.

### Statistics

After identifying and removing outlier values, normality distribution of data was checked using the D’Agostino-Pearson and Shapiro-Wilk tests. Depending on the distribution of the samples (Gaussian or non-Gaussian), different statistical tests were used. For comparisons of one or two samples (one when compared to a hypothetical value), unpaired and paired t-tests were used for Gaussian samples, and Mann Whitney and Wilcoxon signed-rank tests were used for non-Gaussian samples, respectively. When comparing three or more samples, one-way ANOVA and Kruskal-Wallis tests were used for unpaired Gaussian and non-Gaussian samples, respectively, while repeated measurements ANOVA and Friedman tests were used for paired Gaussian and non-Gaussian samples, respectively. For comparing two variables and their interaction, repeated measurements two-way ANOVA test was used for paired Gaussian samples. All statistical analysis was conducted using Prism 7 software (GraphPad Software, Inc., California, USA) and Matlab (The Mathworks Inc., Massachusetts, USA).

## SUPPLEMENTARY FIGURES

**Supplementary Figure 1.**
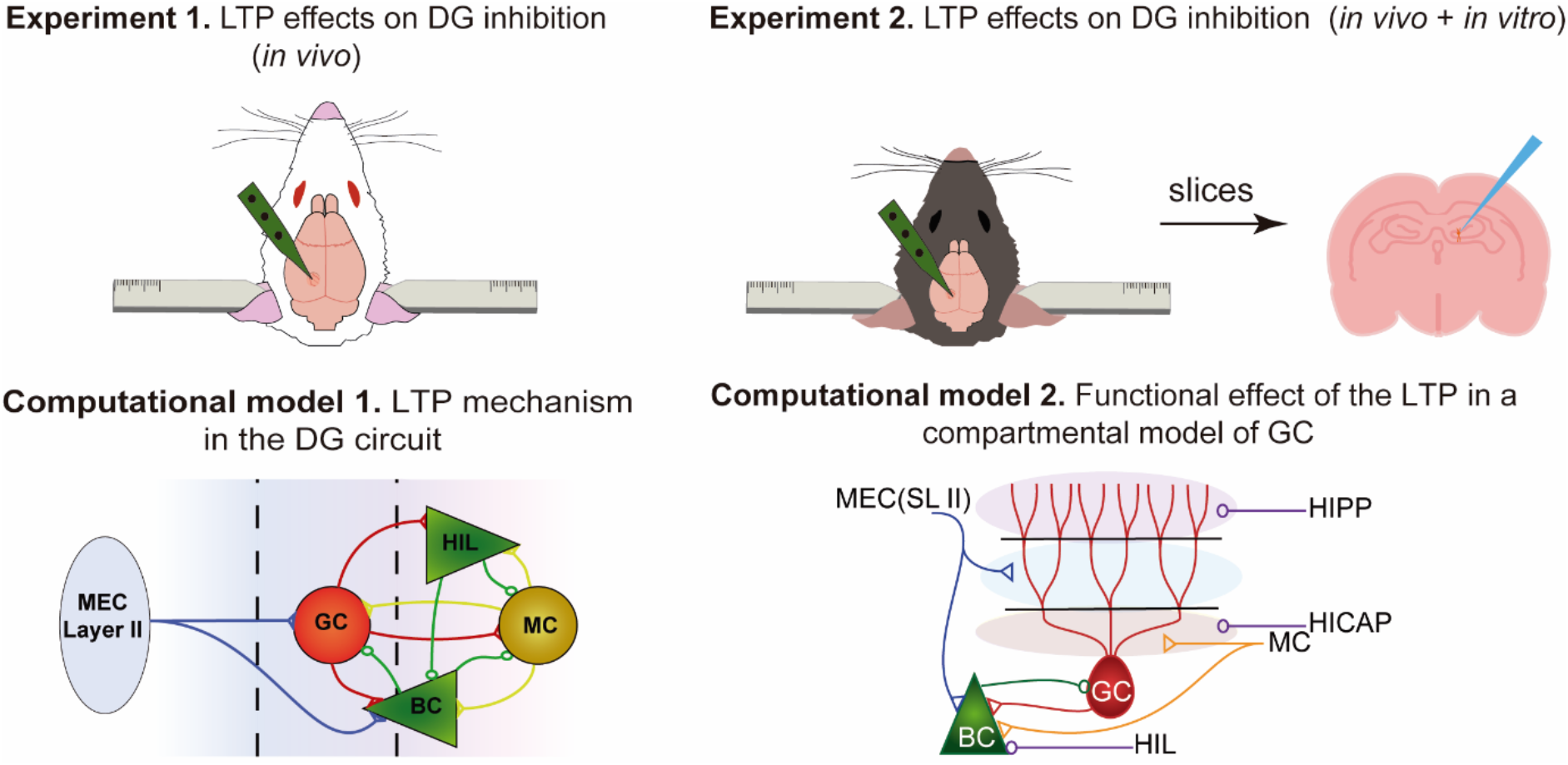
Experimental design. This study employed multichannel electrophysiological recordings *in vivo* on rats (experiment 1) and mice (experiment 2), where long-term potentiation (LTP) was induced in the perforant pathway. The main difference between the two experiments was that mice were sacrificed immediately after LTP induction to perform *in vitro* electrophysiological recordings on slices. To complement the experimental data, we built two computational models: one for a population of dentate gyrus neurons (model 1) and another for a granule cell compartmental circuit (model 2).

**Supplementary Figure 2.**
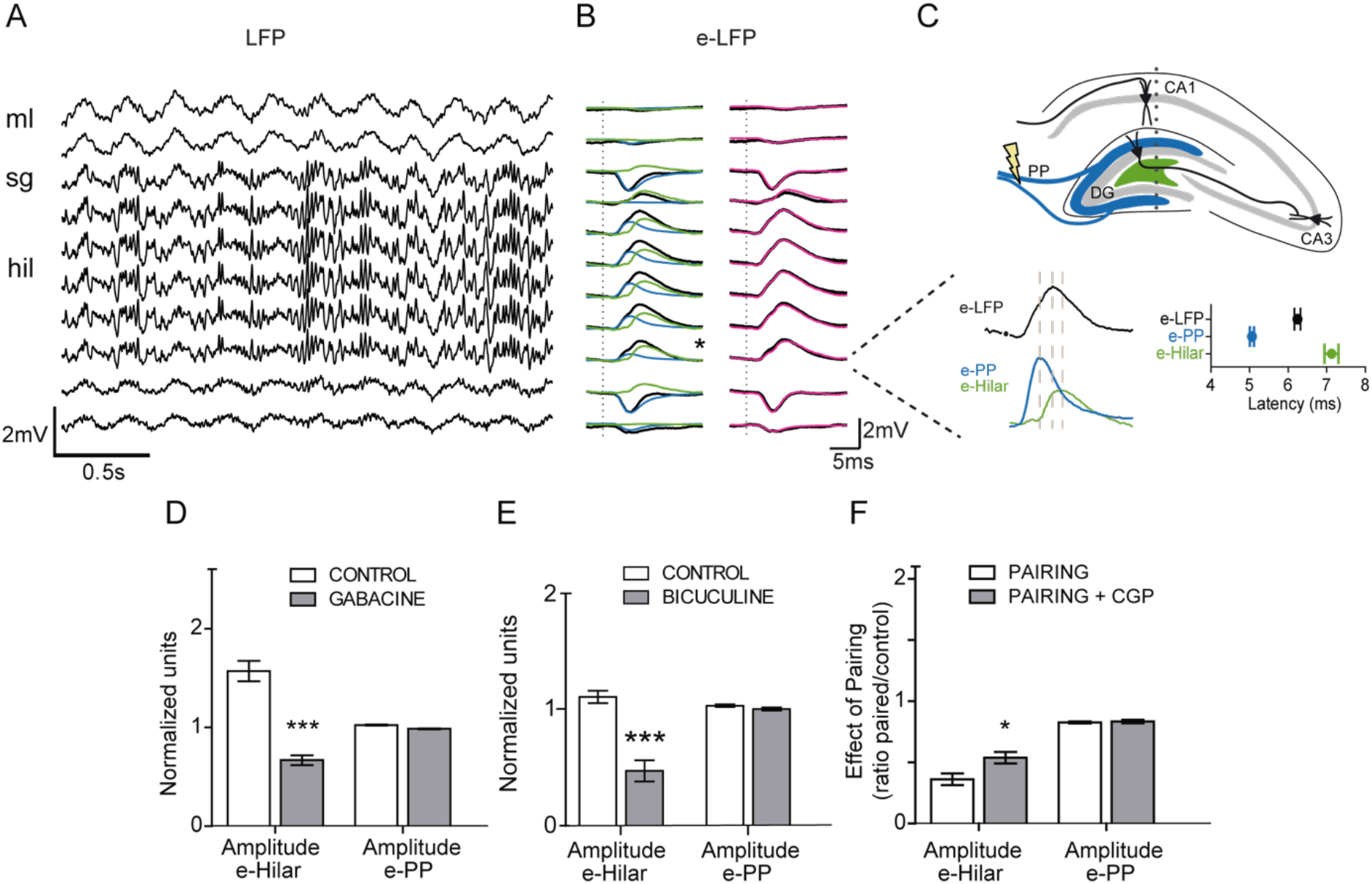
LFP components and pharmacological characterization. **A)** Representative DG recording from 32-channels probe showing spontaneous LFPs acquired in the dorsal hippocampus. The letters on the left mark the anatomical territories of the DG: ml, *molecular layer*; sg, *stratum granular;* hil, *hilus.* **B)** Evoked-LFPs (black) superimposed with the evoked potential recovered in the PP-IC (e-PP) (cyan) and PP-Hilar (e-Hilar) (green) components (left) and with the summation of both components (purple, right). Asterisk marks the potentials enlarged in panel D. **C)** Upper panel. Scheme of a coronal section of the hippocampus and the topographic location of the components. Marked with yellow lightning is the stimulation site. **C)** Lower panel. Same example traces of evoked responses as in B) and group averages of their respective peak latencies (identified with dashed lines). **D, E)** Effects of gabacine and bicuculine, respectively, on e-Hilar and e-PP amplitudes, normalized to the average value in each experiment. Gabacine: Mann Whitney test. U=318, p<0.0001; Bicuculine: Mann Whitney test. U=69, p<0.0001. **F)** Effects of CGP over the pairing pulses on e-Hilar and e-PP amplitudes (measured as the ratio of the value after the pairing by the value before pairing), in control conditions (white) and in the presence of CGP (gray). Paired t-test, t(34)=2.645, p=0.0123. Bars represent mean ± SEM.

**Supplementary Figure 3.**
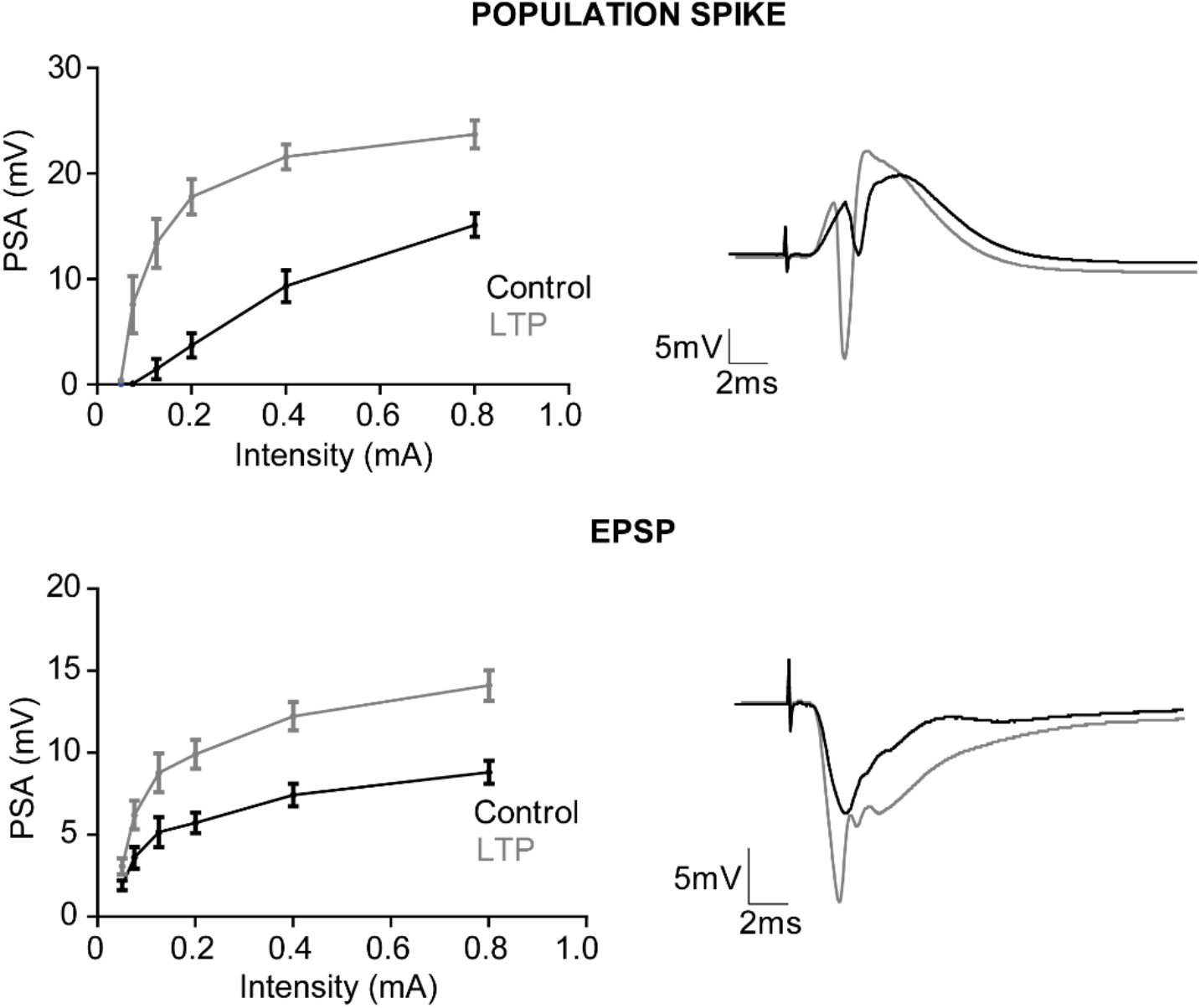
Effects of LTP induction on the evoked-LFPs recorded in DG. Stimulus-response curves showing the amplitude of the PS (upper, left) and the slope of the EPSP (lower, left) evoked by different current intensities (x-axis) before (control) and after LTP induction (LTP). In all panels symbols represent mean ± SEM. Representative example traces of the PS and EPSP before and after LTP are shown on the right. EPSP: 2-way repeated measures ANOVA, Interaction F(_10,80)_=8.86, p<0.0001; Condition F_(2,80)_=227.77, p<0.0001; Intensity F_(5,40)_=21.30, p<0.0001. PS: 2-way repeated measures ANOVA, Interaction F_(10,80)_=17.0, p<0.0001; Condition F_(2,80)_=214.92, p<0.0001; Intensity F_(5,40)_=52.24, p<0.0001.

**Supplementary Figure 4.**
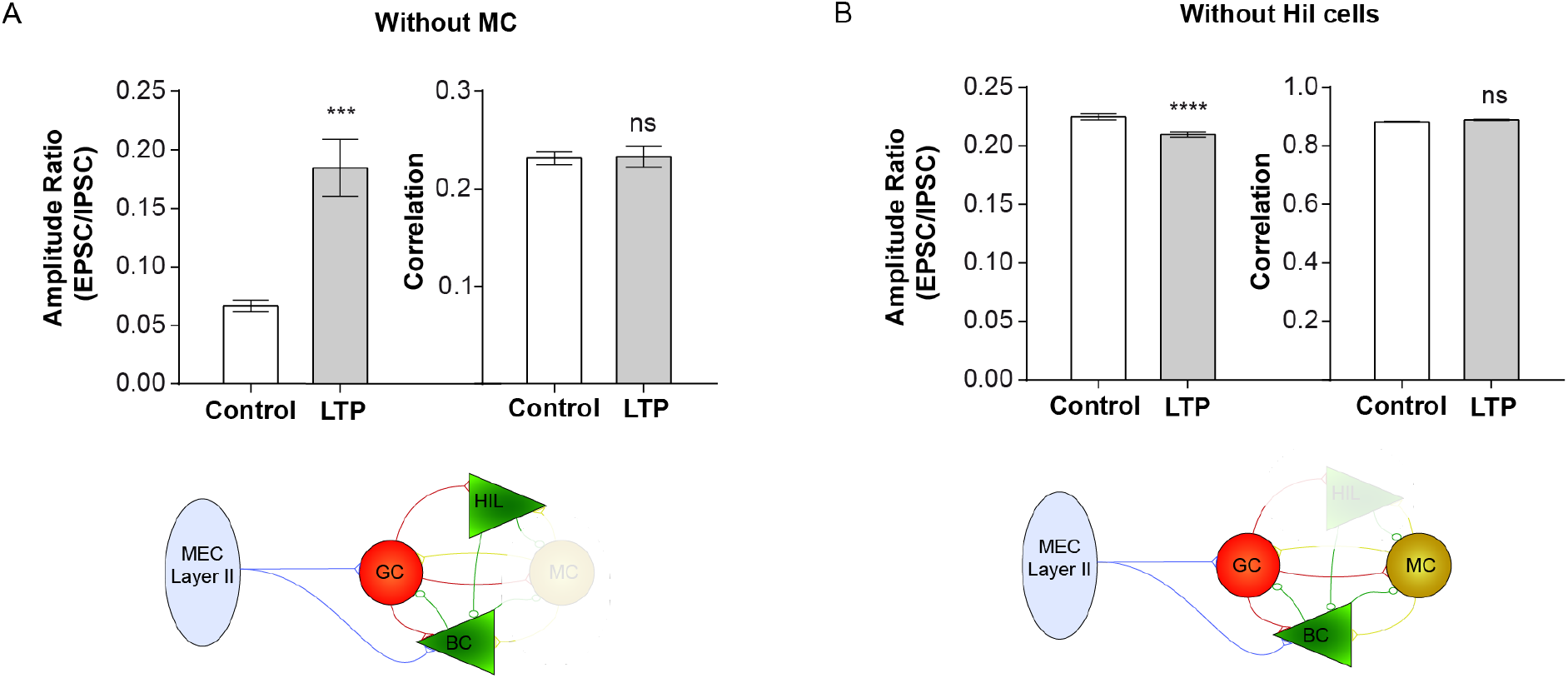
We measured the ratio amplitude and correlation between the excitatory and inhibitory activity onto GCs in two simulation scenarios. A) The first scenario involved the Control circuit and LTP potentiating the MEC input into GC and BC populations, without the MCs. The ratio in this scenario was U=16, p<0.0001. B) The second scenario involved the Control circuit and LTP potentiating the MEC input into GC and BC populations, without the Hil interneurons. The ratio in this scenario was t(38)=4.8, p<0.0001. In all panels, the bar represents the mean ± SEM.

**Supplementary Figure 5.**
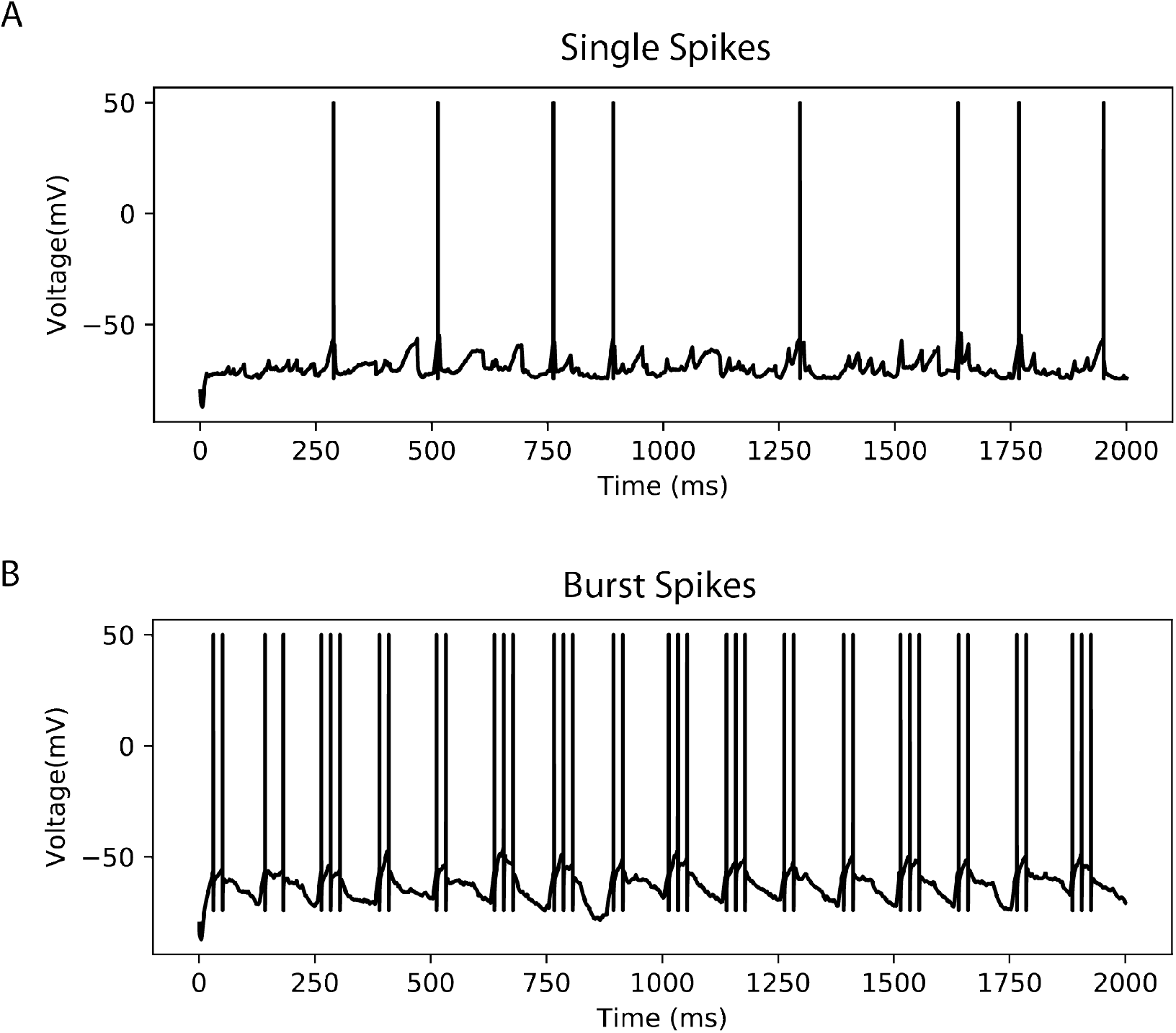
Shown are temporal traces of the membrane potential of a simulated granule cell in computational model 2, with an excitatory input from the MEC at 8 Hz. A) The trace displays a granule cell generating single spikes with baseline inhibitory tone. B) The same granule cell generates burst spikes after decreasing the perisomatic inhibitory tone.

**Supplementary Figure 6.**
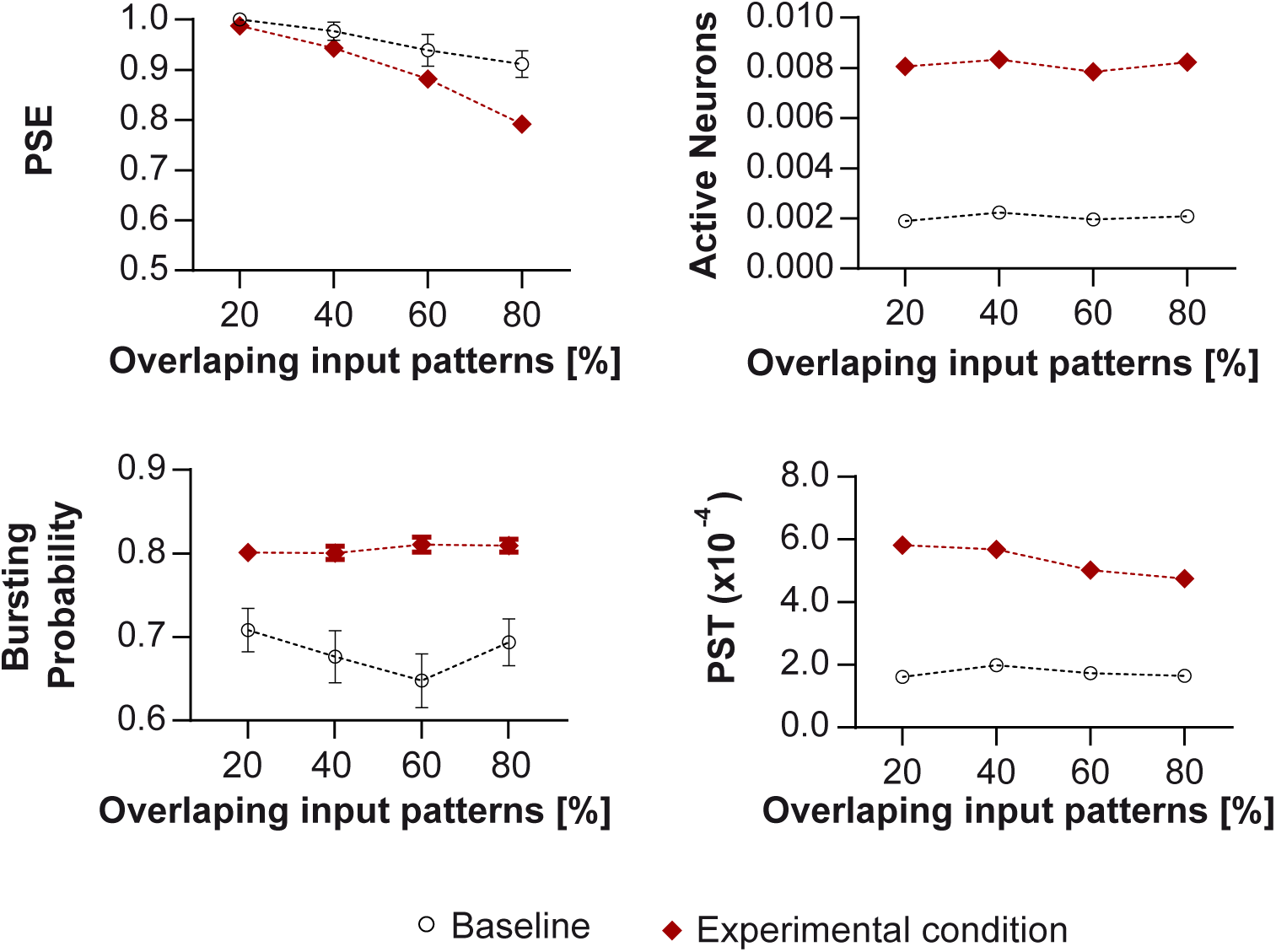
Functional impact of the inhibitory imbalance found experimentally and tested in the granule cell model presented in Figure 4. The experimental case, depicted in red, simulated the effect observed in Figure 2 by introducing two changes, compared to the baseline model, following LTP induction: a 10% reduction in peri-somatic and dendritic inhibitory current, and a 50% reduction in the number of peri-somatic inputs. The results were evaluated by computing the same indicators as in Figure 4. The symbols represent the mean value across 50 analysed pairs, while the error bars indicate the standard error of the mean. Error bars do not appear if their size is smaller than the symbol. The results obtained mimicking the experimental condition qualitatively agree with those of Figure 4.

## Notes

### Competing Interest Statement

The authors have declared no competing interest.

